# Neural and behavioral correlates of edible cannabis-induced poisoning: characterizing a novel preclinical model

**DOI:** 10.1101/2023.03.15.532815

**Authors:** Richard Quansah Amissah, Hakan Kayir, Malik Asfandyaar Talhat, Ahmad Hassan, Yu Gu, Ron Johnson, Karolina Urban, Jibran Y. Khokhar

**Affiliations:** Department of Biomedical Sciences, Ontario Veterinary College, University of Guelph, Guelph, Ontario, Canada; Avicanna Inc., Toronto, Canada; Department of Anatomy and Cell Biology, Schulich School of Medicine and Dentistry, Western University, London, Ontario, Canada

## Abstract

Accidental exposure to Δ^9^-tetrahydrocannabinol (THC)-containing edible cannabis, leading to cannabis poisoning, is common in children and pets; however, the neural mechanisms underlying these poisonings remain unknown. Therefore, we examined the effects of acute edible cannabis-induced poisoning on neural activity and behavior. Adult Sprague-Dawley rats (6 males, 7 females) were implanted with electrodes in the prefrontal cortex (PFC), dorsal hippocampus (dHipp), cingulate cortex (Cg), and nucleus accumbens (NAc). Cannabis poisoning was then induced by exposure to a mixture of Nutella (6 g/kg) and THC-containing cannabis oil (20 mg/kg). Subsequently, cannabis tetrad and neural oscillations were examined 2, 4, 8, and 24 h after THC exposure. In another cohort (16 males, 15 females), we examined the effects of cannabis poisoning on learning and prepulse inhibition, and the serum and brain THC and 11-hydroxy-THC concentrations. Cannabis poisoning resulted in sex differences in brain and serum THC and 11-hydroxy-THC levels over a 24-h period. It also caused gamma power suppression in the Cg, dHipp, and NAc in a sex- and time-dependent manner. Cannabis poisoning also resulted in hypolocomotion, hypothermia, and anti-nociception in a time-dependent manner and impairments in learning and prepulse inhibition. Our results suggest that the impairments in learning and information processing may be due to the decreased gamma power in the dHipp and PFC. Additionally, most of the changes in neural activity and behavior appear 2 hours after ingestion, suggesting that interventions at or before this time might be effective in reversing or reducing the effects of cannabis poisoning.

## INTRODUCTION

Cannabis is mostly consumed through smoking and inhalation [1, 2]; however, the use of edible cannabis, consumed by oral ingestion, has increased in recent times [3-5]. This has resulted in an uptick in accidental cannabis poisoning cases among children and pets [6-9], who mistake edible cannabis for other cannabis-free foods [10]. Moreover, edible cannabis is the most common cannabis product associated with cannabis poisoning in children and pets [11-13], emphasizing the need to investigate the effects of edible cannabis-induced poisoning. Cannabis poisoning in the context of suspected suicide also increased significantly between 2009 and 2021, further emphasizing the need for studying this topic [14]. In children and pets, most clinical signs of cannabis poisoning are neurological [15, 16], occurring due to the interaction between Δ^9^-tetrahydrocannabinol (THC) and type 1 cannabinoid receptors (CB1Rs) expressed in brain regions including the cingulate cortex (Cg), prefrontal cortex (PFC), hippocampus, and nucleus accumbens (NAc). The THC in edible cannabis is metabolized into 11-hydroxy-Δ^9^-tetrahydrocannabinol (11-OH-THC) [17], which might be more potent than THC [18, 19] and cause stronger and longer lasting effects [20]. Oral THC administration also results in high brain THC levels [21], which may contribute to the high incidence of edible cannabis-induced poisoning [22].

Intraperitoneal THC administration causes decreased theta and gamma power in the rat hippocampus, a brain region critical for learning [23], possibly through the inhibition of presynaptic neurotransmitter release [24]. Gamma oscillations are necessary for associative learning, and therefore, their alteration may lead to cognitive deficits [25, 26]. THC vapor administration also resulted in decreased gamma power in the rat dorsal striatum, orbitofrontal cortex, and PFC, regions implicated in the cognitive and psychotic effects of THC [27]. While THC inhalation and injection leads to decreased brain oscillations, few studies have investigated similar effects following edible cannabis consumption. Information processing, evaluated as prepulse inhibition (PPI) in the acoustic startle reflex test (ASR) [28, 29], is impaired in cannabis users [30, 31], and could be used to understand the impact of cannabis poisoning on sensorimotor gating. In rodents, THC administration affects PPI [32-35]. Learning deficits can also be evaluated using the active avoidance task (AAT), and in animal models, acute THC administration causes poor AAT performance [36]. Edible cannabis poisoning may also produce such deficits. While THC exposure via vapor, injection, and gavage induces specific behavioral effects referred to as the cannabis tetrad, which includes catalepsy, hypolocomotion, hypothermia, and anti-nociception [37-39], few studies have evaluated similar behavioral effects using edible cannabis [40, 41]. Sex differences in sensitivity to the effects of THC [14, 42, 43] and its metabolism [44] exist, therefore, it may be worth investigating these differences with respect to edible cannabis poisoning.

This is an important topic given that most patients who report to the emergency unit due to cannabis poisoning report consuming edible cannabis [45]. Understanding the underlying neural mechanisms and the pharmacokinetics of THC via edible consumption could help identify effective treatments for cannabis poisoning. Therefore, the objectives of this study were to investigate the effects of acute high-dose THC via edible administration on neural activity, behavior, and serum and brain THC and 11-OH-THC levels over a 24-h period.

## MATERIALS AND METHODS

### Animals

Forty-four 56 day-old rats (body weight: 150-250 g; males: n=22, females: n=22) were purchased from Charles River for the study. These rats were divided into two cohorts: cohort 1 (males: n=6, females: n=7) and cohort 2 (males: n=16, females: n=15). The rats were singly housed and allowed to habituate for one week. To motivate rats to consume a mix of either Nutella (a brand of sweetened hazelnut cocoa spread) and medium-chain triglyceride (MCT) oil (N-MCT; vehicle for dissolving and diluting THC-containing cannabis oil) or Nutella and THC-containing cannabis oil (N-THC; edible), rats were maintained on 85-90% free-feeding body weight on standard rodent chow for the entire study period, except the first week after arrival and the week following stereotaxic surgery. Rats were maintained on a 12-h light/dark cycle with light on at 07:00 hours. Experiments were performed during the light phase. All procedures were in accordance with guidelines set by the University of Guelph Animal Care Committee and guidelines in the Guide to the Care and Use of Experimental Animals.

### Preclinical model development

On the 7^th^ day after arrival, rats received 3 ml Nutella (density: 1.2 g/ml) for 3 h (Fig. 1A). Food restriction began afterwards and lasted until at least the 13^th^ day. On the 9^th^ and 11^th^ days, rats received N-MCT (Nutella: 6 g/kg, MCT oil: 20 mg/kg) for 10 min. To induce cannabis poisoning, N-THC was prepared similar to N-MCT; however, THC-containing cannabis oil (Five Founders THC Oil [Ontario Cannabis Store]; 30 mg/g THC) was added at 20 mg/kg (adjusted to animal’s body weight). On the 13^th^ day, rats were either given 10-min access to N-THC to induce cannabis poisoning, given N-MCT, or underwent stereotaxic surgery.

**Figure 1.**
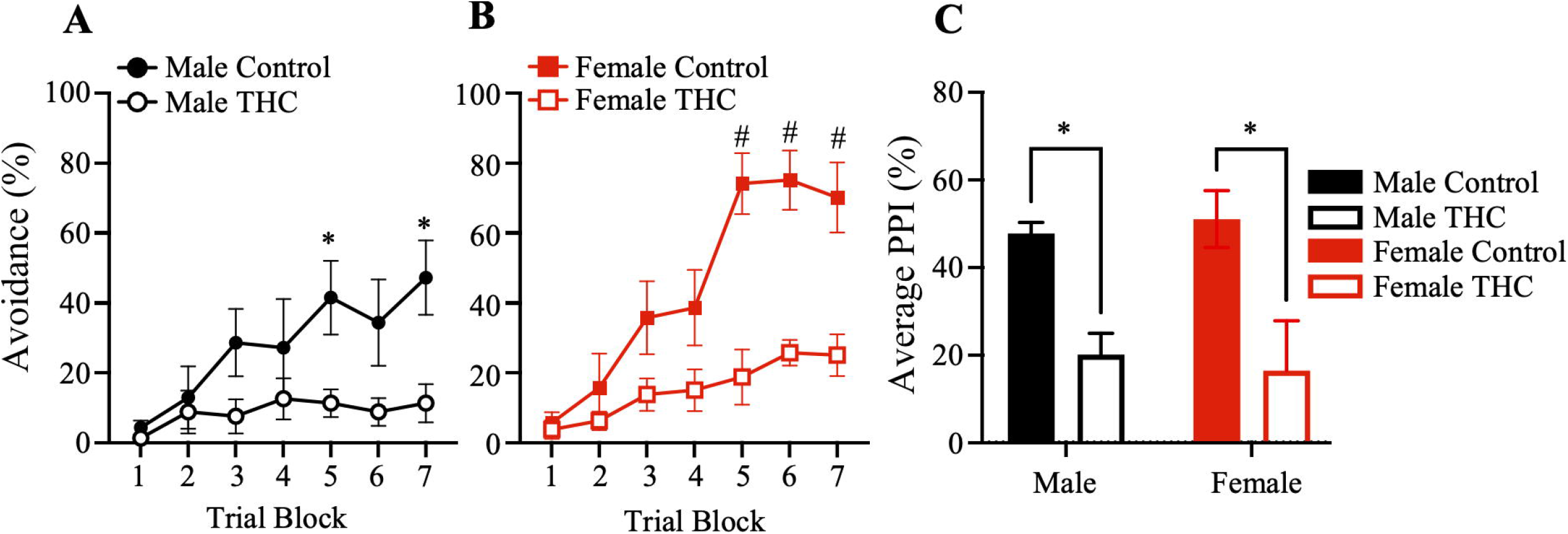
Experiment timeline and details. A. Schematic showing experiments performed each day during the study period. N: Nutella, N-MCT: mix of Nutella and medium-chain triglyceride oil, N-THC: mix of Nutella and THC-containing cannabis oil, C1: cohort 1, C2: cohort 2, ASR: acoustic startle reflex test, AAT: active avoidance test, Ctrl: control rats for cohort 2, Test: test rats for cohort 2, LFP: 15 min local field potential recording. Grey circles: final experiments, Tetrad: tetrad behavior tests, D: day. B. Schematic showing details of final experiments performed over 24 h. Tetrad + LFP: The cannabis tetrad was performed 15 min before LFP recording. Saphenous vein blood draws and brain (for cannabinoid measurements) were collected following AAT/ASR. X: time N-THC was administered, empty circles: experiment performed during the 24-h period, black circles: time post-exposure each experiment was performed during the 24-h period.

### Experiment paradigm

The first rat cohort was used for electrophysiology experiments. These rats underwent cannabis exposure as described above for the development of the preclinical model for cannabis poisoning, except that on day 13, they underwent stereotaxic surgery for multielectrode array implantation (described below). One week after surgery, rats were re-habituated to the N-MCT and tetrad behavior test and the results were used as baseline (time-point 0) for comparison to the tetrad test results from the THC experiment. Afterwards, rats were habituated to the recording chamber, where they were tethered to the headstage for 15 min. The data recorded during headstage habituation were used as baseline local field potential (LFP) activity. During the THC test, performed two days later, rats were given N-THC for 10 min. Subsequently, the cannabis tetrad test followed by LFP activity recording 15 min later was performed 2, 4, 8, and 24 h after N-THC consumption (Fig. 1A). Rats were subsequently sacrificed and their brains retrieved for electrode location verification.

The second rat cohort was used for behavioral (active avoidance test [AAT] and acoustic startle response [ASR]) and pharmacokinetics experiments (Fig. 1A). These rats underwent similar treatment as rats in cohort 1; however, they neither received N-THC immediately nor underwent stereotaxic surgery. On days 13 and 15, rats received N-MCT and underwent habituation and baseline measurements for PPI in the ASR operant box. Subsequently, rats in each sex group were divided into the control (ctrl; male: n=8, female: n=8) and the test (test; male: n=8, female: n=7) groups. On day 17, ctrl rats received N-MCT, while the test rats received N-THC. Two and a half hours later, both groups (ctrl and test) underwent the AAT (Fig. 1B). For the test groups, saphenous blood was also collected 4, 8, and 24 h after N-THC consumption. The brains of the test rats were also retrieved following the 24-h blood collection (Fig. 1B). After 5 days (day 22), the remaining ctrl rats (males: n=8, females: n=8) received N-THC and 3.5 h later, underwent the ASR. Similarly, saphenous blood was collected 4, 8, and 24 h after N-THC consumption (Fig. 1B). Test rat brains were retrieved after the 24-h blood collection.

### Tetrad behavior evaluation

The tetrad tests comprised of tests for hypothermia, catalepsy, hypolocomotion, and analgesia. To test for hypothermia, the rectal temperature of rats was measured using a microprobe thermometer (Model BAT-4, Physitemp Instruments Inc.). Catalepsy was evaluated using an open source automated catalepsy bar as described in a previous study [46]. Hypolocomotion was measured over a 10-min period in an open field box 45 x 45 cm in size. Rats were placed in the middle of the box at the start of the test and their total displacement recorded using Ethovision XT 16.0 video tracking software (Noldus Information Technology). To test for analgesia (thermal pain sensitivity), the tails of rats were placed in the groove on the tail flick analgesia meter (Columbus Instruments, Columbus, OH) containing a radiant heat source and latency to tail flexion was recorded. The tails of rats were removed from the groove after 10 s, if the rat did not do so by itself to prevent tissue damage.

### Two-way active avoidance test

The AAT apparatus comprised of a standard two-way shuttle box (model H10-11R-SC, Coulbourn Instruments, Allentown, PA) placed in a ventilated, isolation chamber (height: 51 cm, width: 53 cm, length: 80 cm; model H10-24, Coulbourn Instruments, Allentown, PA) with a grid floor made of stainless steel bars. A metallic wall partition with a 9 x 9 cm door separated the shuttle box, which contained signal lights, a house light, and an infrared sensor to detect transitions between chambers, into two identical chambers. Scrambled electrical foot shocks were delivered via the grid floor by a precision animal shocker (model H13-15, Coulbourn Instruments, Allentown, PA). The Graphic State software version 5.9 (Coulbourn Instruments, Allentown, PA) was used to program the experimental protocol.

Rats were placed individually in the shuttle box and allowed to habituate for 30 s. They underwent 70 signaled avoidance trials, with intertrial intervals ranging from 15 to 60 s. The trials were divided into 7 blocks of 10 trials each. Each trial consisted of a conditioned stimulus (CS; asynchronously flashing signal and house lights at a frequency of 2 Hz) and an unconditioned stimulus (US; 0.5 mA foot shock). The CS was presented for 10 s and the US was applied during the last 2 s of the CS presentation. Shuttling during the CS prevented the delivery of foot shock (avoidance), while shuttling during the US delivery attenuated the foot shock. The number of avoidance behaviors within each block was detected and expressed as percentages per block.

### Prepulse inhibition of the acoustic startle reflex

The ASR was performed using MedAssociates Acoustic Startle Chambers (MED-ASR-PRO1). Each chamber was soundproof and equipped with a ventilation fan, house light, load cell platform, and two speakers for acoustic stimuli (white noise) delivery. The platform was calibrated by adjusting the load cell amplifier gain, which ranged from -2047 to 2047 arbitrary units, to 200 arbitrary units with a standard weight of 300 g. In the chamber, rats are restrained using a plexiglass cylinder mounted on the platform. A 70-dB white noise was used as background noise during the experiment.

During the baseline session, rats were placed in the plexiglass cylinders in the startle reflex chambers for 15 min, while white noise (background noise) was on. Five acoustic startle sounds were played between 5 and 10 min. In the test session, rats were allowed 5 min to acclimate to the chamber, while the 70 dB background noise was on. Subsequently, five consecutive startle stimuli (120 dB) were presented, followed by 50 trials separated into 10 blocks, and then five more startle stimuli. Each block comprised of five trials in a randomized order: 1. startle stimuli alone, 2. a 73-dB prepulse and startle stimuli, 3. a 76-dB prepulse and startle stimuli, 4. an 82-dB prepulse and startle stimuli, and 5. no stimulus (only background noise). These prepulse intensities were selected because they did not elicit significant startle reflex when applied alone. The startle stimulus lasted for 40 ms, while the prepulse stimulus lasted for 20 ms. The prepulse stimulus was applied 120 ms prior to the onset of the startle stimulus. The intertrial interval ranged from 15 to 30 s. In this study, PPI was defined as a decrease in the magnitude of the startle reflex elicited by a startling stimulus, when it is preceded by a non-startling stimulus (prepulse). The PPI for each prepulse intensity was calculated as follows:

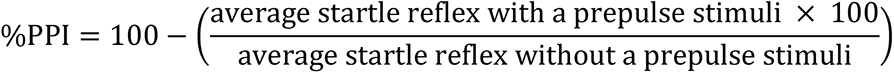

### Serum and Brain THC and 11-OH-THC concentration quantification

#### Materials

Reference standards of 11-OH-THC and THC and their deuterated internal standards 11-OH-THC-D3 and THC-D3 were purchased from Sigma-Aldrich Canada (Oakville, ON). Liquid chromatography mass spectrometry (LC-MS) grade acetonitrile, methanol, and formic acid were purchased from Fisher Scientific (Fisher Chemical Optima™ grade). Ammonium formate was purchased from Sigma (St. Louise, MO, USA) and water was obtained from the Milli-Q system (Millipore, Bedford, MA, USA).

#### Serum Sample Preparation

To extract 11-OH-THC and THC from rat serum, a Captiva enhanced matrix removal lipid (EMR-Lipid) 96-well plate (Agilent, Santa Clara, CA, USA) was used. Briefly, 250 μL of acetonitrile (acidified with 1% formic acid) was added to each well, then 50 μL of rat serum and 20 μL (10 μg/mL) of internal standard solution were added. After the sample passed through under positive pressure at 3 psi, the extraction plate was washed with 150 μL of a mixture of water/acetonitrile (1:4; v:v) and passed through under gradually increasing pressure up to 15 psi. The effluent was evaporated under nitrogen at 40 °C, and the residual was reconstituted with 100 μL of mobile phase for subsequent LC-MS/MS analysis. Calibration Standards (2-1000 ng/mL) and quality controls (3 ng/mL and 800 ng/mL) were prepared on the day of analysis by spiking standard working solutions into blank rat serum.

#### Brain Sample Preparation

The brain tissue samples were removed from -80 °C freezer and immediately placed into a cooled rat brain matrice. A coronal slice between 6 and 7 mm posterior to the frontal tip of brain was cut, weighed and placed into the ice cooled glass tubes for manual homogenization with a glass tissue homogenizer. Acetonitrile (2 mL) and 50 mL internal standard solution (10 μg/mL) were added, and the tissue was homogenized completely. Samples were sonicated in an ice bath for 30 min, and stored overnight at −20 °C. The next day, samples were centrifuged at 1800 rpm at 4 °C for 4 min, and the supernatant were transferred to a new glass tube and evaporated under nitrogen. The tube sidewalls were washed with 250 µL acetonitrile and evaporated again. The samples were reconstituted in 100 µL of the mobile phase, centrifuged for 20 min (15,000 rpm, 4 °C), and the supernatants transferred to the vials. Calibration Standards (1-4000 ng/mL) and quality controls (10 ng/mL and 2000 ng/mL) were prepared on the day of analysis by spiking standard working solutions into the mobile phase.

#### Liquid Chromatography Mass Spectrometry-Mass Spectrometry (LC-MS/MS) method

Serum and brain concentration of 11-OH-THC and THC were determined using an LC-MS/MS method. The LC separation was achieved on a Thermo Scientific Vanquish Flex UHPLC system. Five microliters of sample extracts were injected and separated on an ACQUITY UPLC BEH C18 Column (1.7 µm, 2.1 mm × 50 mm; Waters, Ireland) connected with a VanGuard UPLC BEH C18 Pre-Column (Waters, Ireland). The auto sampler was kept at 4 °C and column temperature was at 35 °C. The mobile phase containing A: 10 mM ammonium formate with 0.1% formic acid aqueous solution, and B: acetonitrile with 0.1% formic acid, was at a flow rate of 400 μL/min under a gradient mode. The gradient conditions were: from 0.1 to 4 min ramp from 40 to 95% of mobile phase B, maintain 95% B for 2 min, then ramp back to 40% B. 11-OH-THC and THC were eluted at 3.5, 4.0 and 4.6 minutes, respectively, with a total run time of 7 min. MS analysis was conducted with a Thermo Q Exactive Focus Orbitrap mass spectrometer equipped with an Ion Max source in positive electrospray ionization (ESI) mode. The source conditions were optimized as spray voltage 3.5 kV, capillary temp 300 °C, and aux gas heater temp 425 °C.

Data were acquired and processed in parallel-reaction monitoring (PRM) mode using Thermo Scientific™ TraceFinder™ software. In this PRM mode, protonated 11-OH-THC ion (*m/z* 331.23) and THC ions (*m/z* 315.23) were selected as precursors, then fragmented in the HCD cell at collision energy of 20 eV for 11-OH-THC and 25 eV for THC. The resulting MS/MS product ions were detected in the Orbitrap at a resolution of 17,500 (FWHM at m/z of 200) with AGC target set at 1e^5^. The most abundant fragments from the MS/MS spectra (*m/z* 313.22 for 11-OH-THC and *m/z* 193.12 for THC) were selected as the quantifying ions. Other specific fragments, m/z 193.12 for 11-OH-THC and m/z 259.17 for THC, were selected as the confirming ions. The resulting chromatograms were extracted and reconstructed with a mass accuracy of 5 ppm for quantification and confirmation. The optimized MS/MS compound parameters are summarized in Table S1.

### Stereotaxic Surgery

After habituation to the N-MCT, rats underwent stereotaxic surgery for electrode implantation, as previously described [47]. A custom-built, 18-channel microelectrode array was constructed using Delrin templates and stainless-steel wires insulated with polyimide tubing. Four regions were targeted bilaterally with eight electrodes: the medial PFC (AP: +3.24 mm, ML: ±0.6 mm, DV: -3.8 mm), Cg (AP: +1.9 mm, ML: ±0.5 mm, DV: -2.8 mm), NAc (AP: +1.9 mm, ML: ±1.2 mm, DV: -6.6 mm), and dHipp (AP: -3.5 mm, ML: ±2.5 mm, DV: -2.6 mm), as per the coordinates found in the Paxinos rat atlas (6^th^ edition) [48]. After the surgery, rats were allowed one week to fully recover before experimentation begun.

### Electrophysiological data acquisition and analysis

LFP data was acquired at a sampling rate of 1 kHz using a TDT RZ10X multichannel acquisition system (Tucker Davis Technologies, FL, USA). The data was filtered between frequencies of 0.1 to 300 Hz. A notch filter was used to remove any 60 Hz frequency noise present in the signal during recording. The recording lasted for 15 min in awake, freely behaving rats in an open field box. LFP analysis was performed using MATLAB (R2020a, The MathWorks^TM^) and routines from the open source Chronux package [49]. The data was preprocessed using detrending and denoising routines in Chronux. Subsequently, the data was low-pass filtered to eliminate frequencies higher than 100 Hz. Afterwards, band pass filters were used to separate the LFP data into its constituent brain rhythms: delta (0.1-4Hz), theta (>4-12 Hz), beta (>12-30 Hz), low gamma (>30-59 Hz), and high gamma (>60-100 Hz). Continuous multispectral power was calculated for each of the brain rhythms in each brain region and coherence between regions was determined.

### Statistical Analysis

The results are presented as means ± standard error of means. The Shapiro-Wilk test was performed to evaluate normality before subsequent statistical tests were performed. The power spectral density for each brain region and the coherence between pairs of brain regions following cannabis poisoning were compared in male and female rats using the two-way repeated measures (RM) analysis of variance (ANOVA). To emphasize within sex differences in male rats following this analysis, the one-way RM ANOVA was used. The two-way RM ANOVA was also performed to compare the means of results obtained during the tetrad behavioral tests, ASR, and serum THC and 11-OH-THC concentrations between male and female rats. Comparison of the brain THC and 11-OH-THC concentrations in male and female rats was performed using the two-tailed unpaired Student’s t-test. The three-way RM ANOVA was performed to compare the means of the percentage avoidance in rats during the AAT, with sex, trial block, and treatment as independent variables and percentage avoidance as the dependent variable. This was followed by a two-way RM ANOVA with treatment and trial block as independent variables based on the lack of sex differences. All ANOVA tests were followed by a post-hoc Bonferroni test to correct for multiple comparisons. Statistical analyses were performed using GraphPad version 6.01 (GraphPad Software Inc., La Jolla CA, USA) and statistical significance was set at p<0.05.

## RESULTS

### Serum and brain THC and 11-OH-THC levels following edible cannabis-induced poisoning

There was a significant main effect of sex (F(1,29)=21.05, p<0.0001), but not time (F(2,58)=2.192, p=0.1208), after N-THC consumption and a sex x time interaction (F(2,58)=0.8209, p=0.4451), on serum THC concentrations. In males, the THC concentration at all time-points were similar (Fig. 2A). In females, the THC concentration at the 4-h time-point was higher than that at the 24-h (p=0.0463) time-point. Moreover, the THC concentration in females was higher than that in males at the 4-h time-point (p=0.0019), but not at the other time-points. There were significant main effects of sex (F(1,29)=86.00, p<0.0001), time (F(2,58)=23.86, p<0.0001), and their interaction (F(2,58)=21.25, p<0.0001) on the serum 11-OH-THC concentration. In males, there were no time-related differences in 11-OH-THC concentration, while in females, the 11-OH-THC concentration at the 4-h time-point was higher than those at the 8-h (p=0.0220) and 24-h (p<0.0001) time-points (Fig. 2B). Additionally, the 11-OH-THC concentration in females was higher than that in males at the 4-h (p<0.0001) and 8-h (p<0.0001) time-points, but not the 24-h time-point. Comparison using the t-test revealed higher brain THC and 11-OH-THC levels in females (p=0.0159 and p=0.0085, respectively) compared to males at the 24-h time-point (Fig. 2C and D).

**Figure 2.**
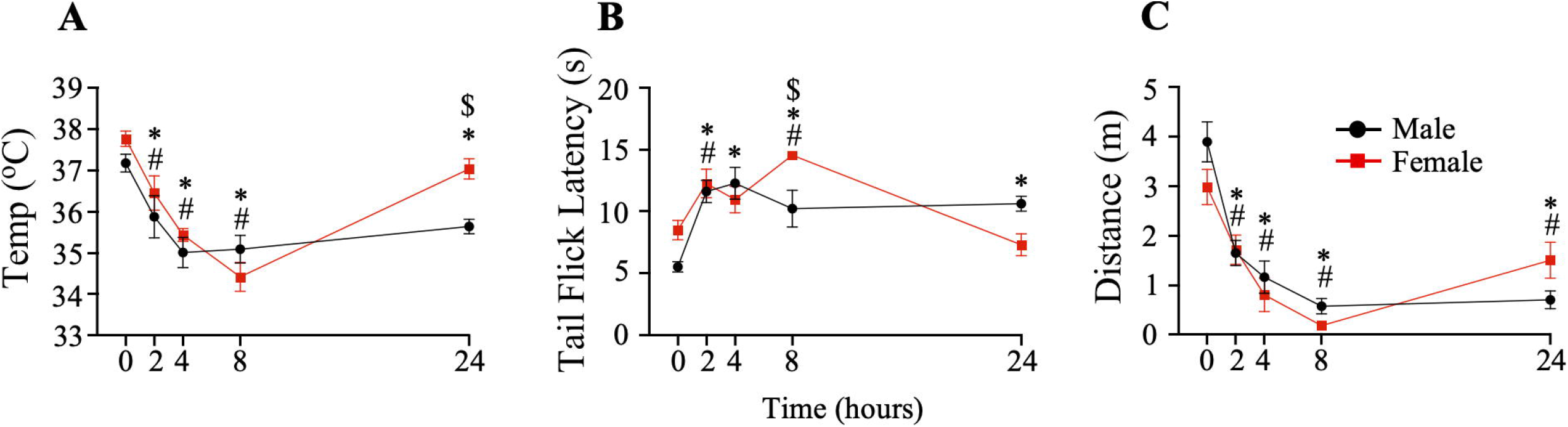
Concentrations of THC and 11-OH-THC in the serum and brain of rats following edible cannabis-induced poisoning. A and B. Serum THC and 11-OH-THC concentrations in males and females at the 4-, 8-, and 24-h time-points. The serum THC and 11-OH-THC concentrations in females were generally higher than those in males. C and D. Brain THC and 11-OH-THC levels in males and females at the 24-h time-point. The brain THC and 11-OH-THC levels in females were higher than those in males. Filled black circles: Male THC/11-OH-THC, filled red squares: female THC/11-OH-THC. $: comparison between males and females at different time-points with p<0.05. #: comparison of concentration at other time-points with that at the 4-h time-point in females with p<0.05. *: comparison of concentration of THC/11-OH-THC in the brain between males and females with p<0.05.

### Effects of edible cannabis-induced poisoning on gamma oscillations

Although N-THC consumption had some effect on delta, theta, and beta oscillations, the low and high gamma oscillations were the most consistently affected, consistent with previous findings with THC and cannabis [27, 50, 51]. Therefore, only the results for the effects of acute high-dose THC via edible administration on PFC, dHipp, Cg, and NAc gamma oscillations at the different time-points (Fig. 3) will be described.

**Figure 3.**
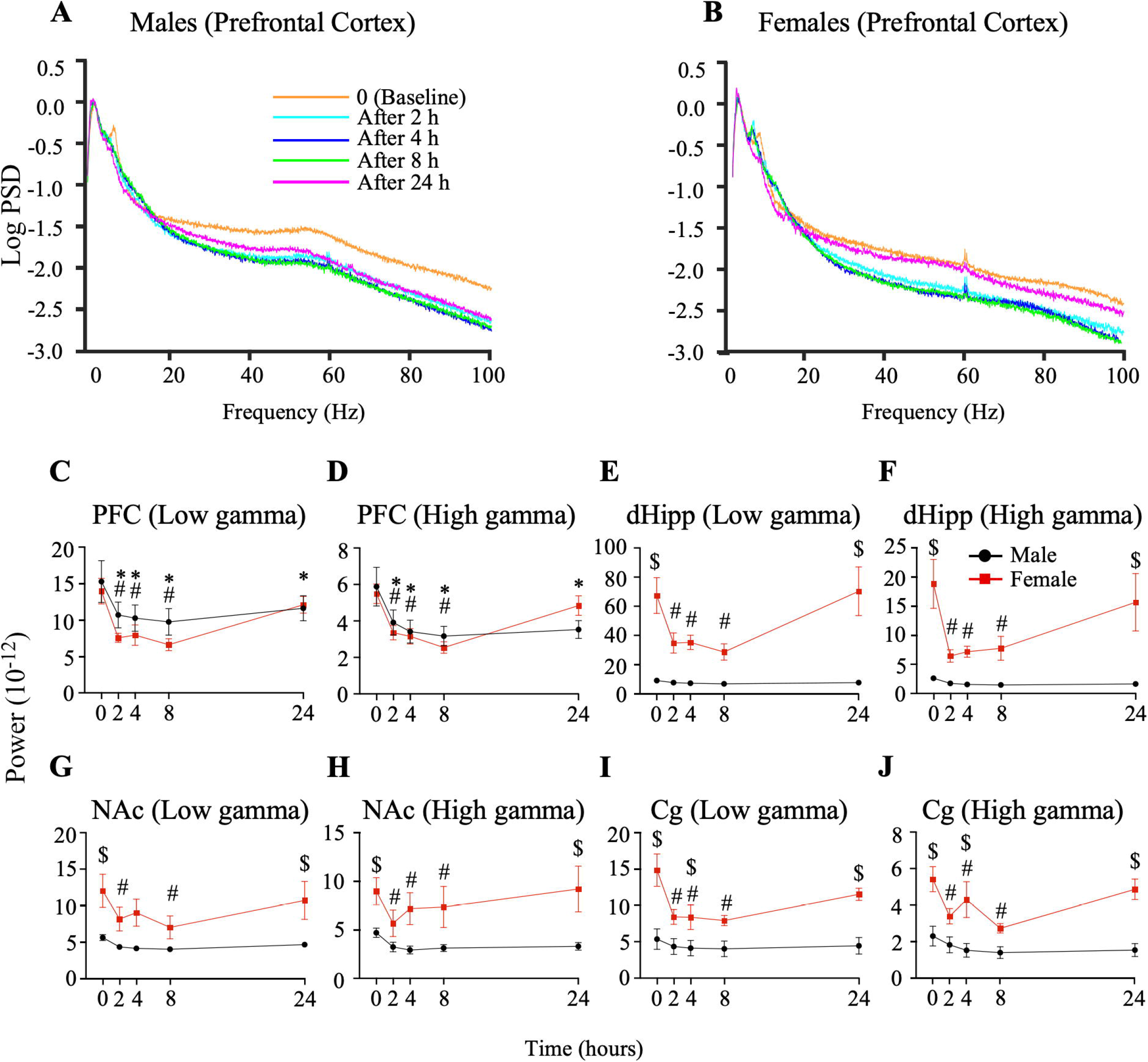
Power spectral density (PSD) plot for male and female rats after edible cannabis-induced poisoning. Representative log-transformed prefrontal cortex (PFC) PSD in male (A) and female rats (B) at each time-point. Orange plot: Baseline, cyan plot: 2-h time-point, blue plot: 4-h time-point, green plot: 8-h time-point, and pink plot: 24-h time-point. C and D. Low and high gamma power, respectively, in the PFC between males and females at different time-points. The low and high gamma power in the PFC of male and female rats were generally not different at the targeted time-points. E and F. Low and high gamma power, respectively, in the dorsal hippocampus (dHipp) of males and female rats at different time-points. G and H. Low and high gamma power, respectively, in the nucleus accumbens (NAc) of males and female rats at different time-points. I and J. Low and high gamma power, respectively, in the cingulate cortex (Cg) between males and females at different time-points. In general, the low and high gamma power in the dHipp, Cg, and NAc of females were higher than those in males (E – J). $: comparison of time-points between sexes with p<0.05, *: comparison of power at other time-points with that at baseline in males with p<0.05, #: comparison of power at other time-points with that at baseline in females with p<0.05.

In the PFC, while the main effects of sex (F(1,24)=0.7462, p=0.3962) and its interaction with time (F(4,96)=1.946, p=0.1093) on the low gamma (LG) power were not significant, that for time (F(4,96)=17.59, p=0.0001) was significant. Moreover, the LG power was similar in males and females at all time-points (Fig. 3A-D). However, in males the LG power at the 2-h (p=0.0035), 4-h (p=0.0012), 8-h (p=0.0003), and 24-h (p=0.028) time-points were lower than that at baseline. In females, with the exception of the 24-h LG power, the 2-h (p<0.0001), 4-h (p<0.0001), and 8-h (p<0.0001) LG power was lower than that at baseline. The main effect of sex on high gamma (HG) power in the PFC (F(1,24)=0.02193, p=0.8835) was not significant; however, that for the time (F(4,96)=23.42, p<0.0001) and its interaction with sex (F(4,96)=3.169, p=0.0171) were significant. Additionally, the PFC HG power at baseline and at all time-points were similar in males and females (Fig. 3A-D). The HG power at the 2-h (p=0.002), 4-h (p<0.0001), 8-h (p<0.0001), and 24-h (p<0.0001) time-points were lower than that at baseline in males. In females, the HG power at the 2-h (p<0.0001), 4-h (p<0.0001), and 8-h (p<0.0001) time-points, but not the 24-h time-point, was lower than that at baseline.

Significant main effects of sex (F(1,24)=24.40, p<0.0001), time (F(4, 96)=5.540, p=0.0005), and their interaction (F(4,96)=4.897, p=0.0012) on dHipp LG power were observed. Moreover, while the LG power at baseline (p<0.0001) and the 24-h time-point (p<0.0001) were higher in females, it was similar between sexes at the other time-points (Fig. 3E). Furthermore, the LG power at the other time-points were not different from that at baseline in males. In females, however, with the exception of the 24-h LG power, the LG power at the 2-h (p=0.0007), 4-h (p=0.0009), and 8-h (p<0.0001) time-points were lower than baseline. Similarly, the main effects of sex (F(1,24)=14.38, p=0.0009), time (F(4,96)=5.5, p=0.0005), and their interaction (F(4,96)=4.294, p=0.0031) on dHipp HG power were significant. Even though the HG power in both sexes were similar at the 2-h, 4-h, and 8-h time-points, those at baseline (p<0.0001) and the 24-h (p=0.0002) time-point were higher in females (Fig. 3F). In females, but not males, the baseline HG power was higher than those at the 2-h (p<0.0001), 4-h (p<0.0001), and 8-h (p=0.0001) time-points. The one-way RM ANOVA revealed that in males, the baseline LG power was higher than that at the 4-h (p=0.0118) and 8-h (p=0.0044) time-points, but not different from those at the 2-h and 24-h time-points (Fig. S1A), while the HG power at the 2-h (p=0.0011), 4-h (p<0.0001), 8-h (p<0.0001), and 24-h (p<0.0001) were lower than that at baseline (Fig. S1B).

The main effects of sex (F(1,24)=8.994, p=0.0062) and time (F(4,96)=4.122, p=0.0041), but not their interaction (F(4,96)=1.479, p=0.2151), on NAc LG power were significant. The LG power at baseline (p=0.0109) and the 24-h (p=0.0071) time-point were higher in females than in males, but not different at the 2-h, 4-h, and 8-h time-points (Fig. 3G). In males, the LG power at all time-points were similar; however, in females, the LG power at the 2-h (p=0.0130) and 8-h (p=0.0009) time-points were higher than that at baseline, while those at the 4-h and 24-h time-points were not. The sex (F(1,24)=5.544, p=0.0271) and time (F(4,96)=7.510, p<0.0001), but not their interaction (F(4,96)=1.578, p=0.1867), had significant main effects on the NAc HG power. Females had higher HG power at baseline (p=0.0357) and the 24-h (p=0.0384) time-point than males, but similar HG power to males at the other time-points (Fig. 3H). Additionally, the HG power at baseline in males was similar to those at other time-points. In females, the HG power at the 2-h (p=0.0002), 4-h (p<0.0001), and 8-h (p=0.0002) time-points, but not the 24-h time-point, were higher than that at baseline. In the male NAc, the one-way RM ANOVA revealed that, with the exception of the 24-h LG power, the LG power at the 2-h (p=0.0158), 4-h (p=0.0028), and 8-h (p=0.0018) time-points were lower than that at baseline (Fig. S1C). The baseline HG power in males was not different from that at the 2-h time-point but higher than those at the 4-h (p=0.0002), 8-h (p=0.008), and 24-h (p=0.0002) time-points (Fig. S1D).

Significant main effects of sex (F(1,24)=14.69, p=0.0008), time (F(4,96)=8.707, p<0.0001), and their interaction (F(4,96)=4.506, p=0.0023) on Cg LG power were observed. Compared to males, the LG power in females were higher at baseline (p<0.0001) and the 24-h time-point (p=0.0002), but not different at the remaining time-points (Fig. 3I). Moreover, while the LG power at baseline was similar to that at the other time-points in males, the LG power at baseline was higher than that at the 2-h (p<0.0001), 4-h (p<0.0001), and 8-h (p<0.0001) time-points, but not the 24-h time-point, in females. Similarly, there were significant main effects of sex (F(1,24)=14.92, p=0.0007), time (F(4,96)=7.109, p<0.0001), and their interaction (F(4,96)=3.107, p=0.0190) on the Cg HG power. Moreover, while the HG power at the 2-h and 8-h time-points were not different in both sexes, it was lower in males compared to females at baseline (p=0.0004) and at the 4-h (p=0.0048) and 24-h (p=0.0001) time-points (Fig. 3J). Males had similar HG power at all time-points when compared to baseline, while females had HG power at baseline that was higher than that at the 2-h (p=0.0006), 4-h (p=0.0429), and 8-h (p<0.0001) time-points, but not the 24-h time-point. The one-way RM ANOVA revealed that, in males, while the Cg LG power at baseline was not different from that at the other time-points (Fig. S1E), the HG power at baseline was not different from that at the 2-h time-point, but higher than those at the 4-h (p=0.0137), 8-h (p=0.0344), and 24-h (p=0.0320) time-points (Fig. S1F).

The coherence between pairs of brain regions within the gamma frequency ranges was also evaluated. However, unlike the power spectral density analysis, the results (Fig. S2) were inconsistent and will not be described.

### Effects of edible cannabis-induced poisoning on tetrad behavior

While we evaluated the four cannabis tetrad behaviors [39], there were no observable cataleptic effects of cannabis poisoning in both sexes; therefore, results will only be presented for hypolocomotion, hypothermia, and anti-nociception.

There were significant main effects of time (F(4,44)=26.91, p<0.0001) and sex x time (F(4,44)=3.295, p=0.0191) on rectal body temperature, with the cannabis exposed animals showing a hypothermia phenotype. Moreover, in males, while the body temperatures at the 2-h (p=0.0143), 4-h (p<0.0001), 8-h (p<0.0001), and 24-h (p=0.0029) time-points were decreased compared to that at baseline, in females, the rectal temperatures were decreased at the 2-h (p=0.0064), 4-h (p<0.0001), and 8-h (p<0.0001) time-points, but not the 24-h time-point (Fig. 4A). Additionally, the 24-h time-point rectal temperature of females was higher than that of males (p=0.014).

**Figure 4.**
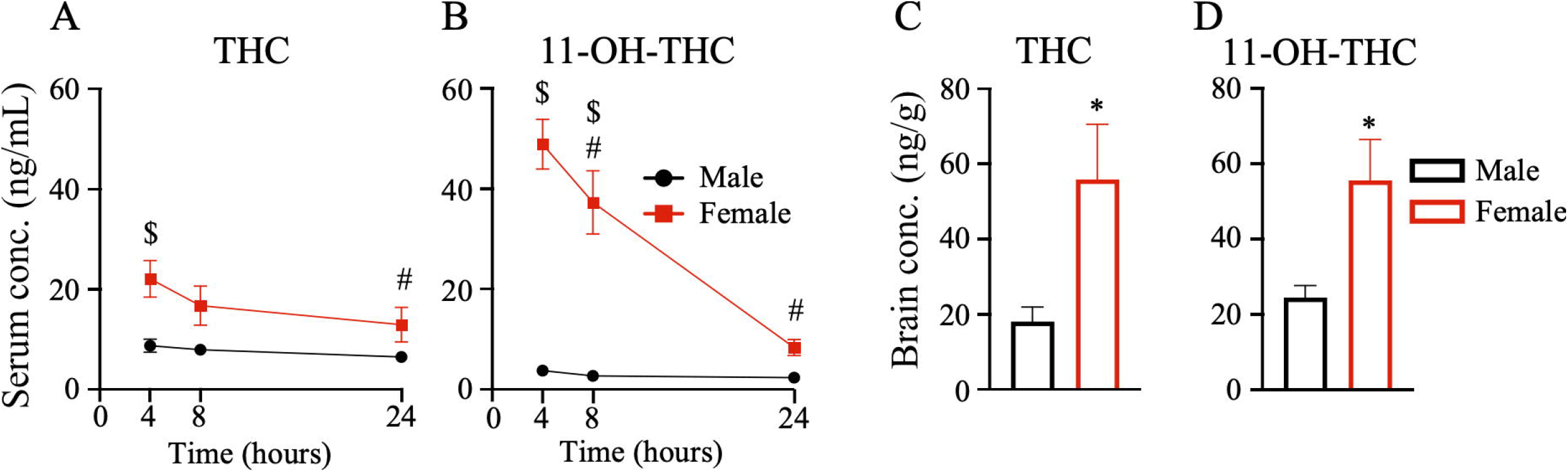
Edible cannabis-induced effects on body temperature, anti-nociception and locomotion in male and female rats. A. Graph comparing the rectal temperatures of males and females measured at the time-points of interest following edible cannabis-induced poisoning. While the rectal temperature at the different time points were different when compared to baseline within male rats and within female rats, the rectal temperature was not different at all time points when male rats were compared to female rats. B. Graph comparing the latency to tail flick of males and females at the time-points of interest following edible cannabis-induced poisoning. While the tail flick latency (anti-nociception) at the different time points when compared to baseline were different within male rats and within female rats, the tail flick latency was not different at all time points when male rats were compared to female rats. C. Graph comparing the total distance traveled by males and females in the open field box at the time-points of interest following edible cannabis-induced poisoning. While the distance traveled in the open field box at the different time points when compared to baseline were different within male rats and within female rats, the distance traveled was not different at all time points when male rats were compared to female rats. $: comparison of outcome at different time-points between sexes with p<0.05, *: comparison of outcome at other time-points with that at baseline in males with p<0.05, #: comparison of outcome at other time-points with that at baseline in females with p<0.05.

There were significant main effects of time (F(4,44)=11.92, p<0.0001) and sex x time interaction (F(4,44)=5.517, p=0.001) on edible cannabis-induced anti-nociception. The latency to tail flick in males was longer at the 2-h (p=0.0003), 4-h (p<0.0001), 8-h (p=0.0067), and 24-h (p=0.003) time-points compared to baseline. In females, the tail flick latency was longer at the 2-h (p=0.0209) and 8-h (p<0.0001) time-points but not at the 4-h and 24-h time-points (Fig. 4B). Moreover, the tail flick latency of females was longer compared to males after 8 h (p=0.0091) and tended to be shorter than that in males at the 24-h (p=0.0743) time-point.

There was a significant main effect of time on displacement (distance traveled) (F(4,44)=32.90, p<0.0001), with animals showing hypolocomotion after edible cannabis exposure. However, the main effect of sex (F(1,11)=0.5725, p=0.4652) was not significant, while that for the sex x time interaction (F(4,44)=2.453, p=0.0598) trended toward significance. Additionally, compared to displacement at baseline, the displacement of both males and females at the 2-h (male: p<0.0001, female: p=0.0092), 4-h (male: p<0.0001, female: p<0.0001), 8-h (male: p<0.0001, female: p<0.0001), and 24-h (male: p<0.0001, female: p=0.0019) time-points were decreased (Fig. 4C).

### Effects of edible cannabis-induced poisoning on active avoidance learning and prepulse inhibition

In the AAT, the three-way ANOVA revealed significant main effects of trial block (F(4.322,109.5)=30.14, p<0.0001), treatment (control vs THC groups; F(1,26)=15.94, p=0.0005), trial block x treatment interaction (F(6,152)=10.29, p<0.0001), and trial block x sex interaction (F(6,152)=4.318, p=0.0005) on percentage avoidance. There were no significant main effects of sex, sex x treatment interaction, or trial block x treatment x sex interaction on percent avoidance. We subsequently performed a two-way ANOVA in males which revealed significant main effects of trial block (F(6,78)=7.037, p<0.0001), treatment (1, 13)=4.701, p=0.0493), and their interaction (F(6,78)=3.396, p=0.0050) on the percentage avoidance (Fig. 5A). The percentage avoidance in control rats during trial blocks 5 (p=0.0474) and 7 (p=0.0098) was higher than that in THC rats but not different during trial blocks 1, 2, 3, 4, and 6. In females, the two-way ANOVA revealed significant main effects of trial block (F(6,74)=27.47, p<0.0001), treatment (F(1,13)=12.79, p=0.0034), and their interaction (F(6,74)=7.552, p<0.0001) on the percentage avoidance. While the percentage avoidance in female control rats during trial blocks 1, 2, 3, and 4 was not different from that for female THC rats, it was higher than that of female THC-exposed rats during trial blocks 5 (p<0.0001), 6 (p=0.0004), and 7 (p=0.0001) (Fig. 5B).

**Figure 5.**
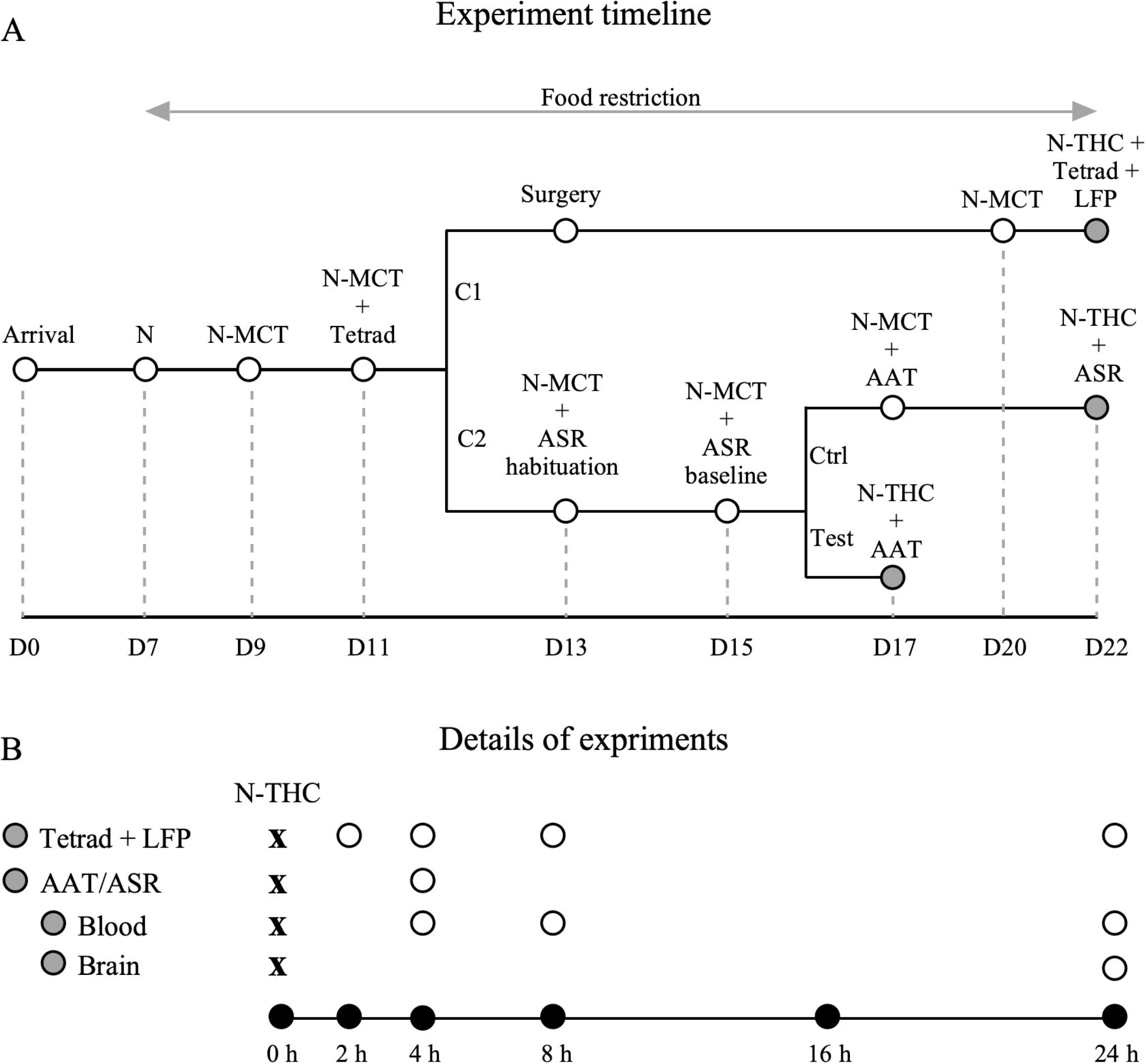
Both active avoidance and prepulse inhibition are disrupted by edible cannabis poisoning. A. Graph comparing the percentage avoidance among male control and THC rats. The percentage avoidance of male control rats was higher than that of male THC rats during trial blocks 5 and 7. Filled black circle: male control rats, empty black circle: male THC rats. B. Graph comparing the percentage avoidance among female control and THC rats. The percentage avoidance of female control rats was higher than that of female THC rats during trial blocks 5, 6, and 7. Filled red square: female control rats, empty red square: female THC rats. *: comparison of percentage avoidance for different trials between male control rats and male THC rats with p<0.05, #: comparison of percentage avoidance for different trials between female control rats and female THC rats with p<0.05. C. Graph comparing average percentage PPI among male and female control and THC rats. The average percentage PPI in both male and female THC rats was lower than those of their corresponding control rats. Filled black bar: male control rats, empty black bar: male THC rats, filled red bar: female control rats, empty red bar: female THC rats. *: comparison between control and THC rats with p<0.05.

During the two-way ANOVA analysis of the percentage PPI, no difference in the effect of sound intensity on the percentage PPI was observed in both sexes (results not shown); the percentage PPI for the three intensities were therefore averaged and analyzed. While the main effects of sex (F(1,14)=0.001341, p=0.9713) and sex x treatment interaction (F(1,14)=0.3888, p=0.5429) on percentage PPI were not significant, that for treatment was significant (F(1,14)=13.92, p=0.0022). Moreover, in both sexes, the averaged percentage PPI for control rats were higher than that for THC exposed rats (male: p=0.0209, female: p=0.0037) (Fig. 5C).

## DISCUSSION

In this study, we established and characterized a rat model of high-dose edible cannabis poisoning. We observed higher serum and brain THC and 11-OH-THC levels in female rats compared to male rats after cannabis poisoning despite exposure to the same N-THC doses. The study also revealed sex- and time-dependent decreases in dHipp, Cg, and NAc gamma power over a 24-h period starting 2 h after cannabis poisoning. Time-dependent changes in the cannabinoid tetrad behaviors and active avoidance and PPI disruptions were also observed.

The observed sex difference in serum THC levels could be partly due to the rapid redistribution of the lipophilic THC to fatty tissues to a greater extent in males [52-54], who have more body fat than females [55]. Similar to a previous study [56], we found sex differences in serum 11-OH-THC levels, suggesting differences in THC metabolism, possibly due to the sexually dimorphic expression of cytochrome P450 (CYP) enzymes [57, 58], resulting in THC being preferentially metabolized into 11-nor-9-carboxy-Δ^9^-THC (11-COOH-THC) in males and into 11-OH-THC in females [59-61], leading to higher serum 11-OH-THC levels in females. We also observed lower brain THC and 11-OH-THC levels in males compared to females, which may be related to the lower serum levels of the parent compound in males. Further, male rat brains may be protected against THC through increased expression of blood brain barrier proteins (including claudin-5) [60, 61]. The rat brain also expresses CYP enzymes that can metabolize THC [62], suggesting that microsomes in female rats may metabolize more THC into 11-OH-THC, resulting in higher brain 11-OH-THC levels [44]. In general, the serum and brain 11-OH-THC levels decreased gradually over 24 h; while this may be due to its excretion [63], it could be because the 11-OH-THC is rapidly oxidized into 11-COOH-THC [64]. The higher brain and serum levels of THC and 11-OH-THC in female rats compared to male rats may also explain the sex differences in the effects of cannabis observed in humans, with female sex being associated with a greater likelihood of cannabis poisoning-related suspected suicide [14, 42, 43].

The local field potential power reflects the extent of neural synchrony, with higher power reflecting higher synchrony and vice versa [65]. Gamma power and synchrony is modulated by parvalbumin-expressing (PV) interneurons [66], which are modulated by cholecystokinin-expressing (CCK) interneurons [67], the only CB1R-expressing interneuron in the cortex and hippocampus [68-70]. THC binding to CCK interneuron CB1Rs affect PV interneuron activity, resulting in a disruption in gamma oscillations, and a subsequent gamma power decrease. This may explain the decreased gamma power observed following edible cannabis-induced poisoning in our rats [27, 50, 71, 72]. That these decreases in gamma power were observed in the dHipp, Cg, NAc, and PFC, is not surprising since these regions have high CB1R densities. In our study, significant edible cannabis-induced poisoning effects on gamma power began around the 2-h mark, which is consistent with the finding that the subjective behavioral effects of edible cannabis peak between 1.5 and 3 h after ingestion [73, 74], as well as our pharmacokinetic findings. We also observed sex differences in NAc, Cg, and dHipp gamma power. This may be partly attributable to the high estrogen levels in female rats [75, 76], which increases the binding of estrogen to estrogen β receptors expressed by PV interneurons, leading to increased firing, greater inhibition, and increased gamma activity [77, 78]. This may explain the higher gamma power in female rats, which may also underlie the quicker recovery from the THC-induced suppression in females, where the 24-h time-points no longer showed THC effects, in addition to potential pharmacokinetic differences or tolerance.

The similarity in effects of THC on anti-nociception (tail flick latency) in males and females observed in our study was also reported in rats following THC vapor administration [40]. One of the most cited reasons for medical cannabis use is pain relief [79, 80]. The periaqueductal gray, a brain region that contains CB1R-expressing somatodendritic structures and presynaptic terminals [81-83], is involved in pain modulation [84], and may play a role in the decreased pain sensitivity we observed following edible cannabis-induced poisoning. Interestingly, the tail flick latency in female rats had returned to baseline levels, unlike that in male rats, 24 h after acute high-dose THC administration. We also observed edible cannabis-induced hypothermia in both males and females, which may be due to the interaction between THC and the preoptic area of the hypothalamus, a region involved in temperature regulation [85] and known to contain high densities of CB1Rs [86-88]. In female rats, both anti-nociception and body temperature had returned to baseline levels by the 24-h time-point, which could be due to the development of tolerance to the higher serum and brain THC and 11-OH-THC levels or the differential time-course of effects between the parent compound and metabolite. While no sex-differences in hypolocomotion was observed in our study, similar to the findings of a previous study [89], the hypolocomotion, which was also previously reported [90, 91], may be due to the interaction between THC, 11-OH-THC, and CB1Rs on cerebellar basket cells [92]. Interestingly, the edible cannabis-induced tetrad effects began around the 2-h time-point, similar to the observed neural effects of cannabis poisoning in this study, which coincides with the time-point that cognitive and psychomotor deficits were observed in humans following edible cannabis ingestion [74]. This suggests that interventions to reverse the effects of edible cannabis-induced poisoning should target the 2-h time window.

The AAT and ASR were performed just before the 4-h time-point because the effects of edible cannabis on cognitive performance in humans were observed between 2 and 5 h post-ingestion [74]. We observed decreased hippocampal gamma power, which is necessary for learning the association between stimuli [26, 93, 94]. This could explain the inability of N-THC-treated rats to avoid the foot shock during the AAT. Our finding is consistent with those of previous studies that found learning deficits in cannabis users [95] and THC-treated rats [36]. While low dose THC (0.3 – 3 mg/kg) administration had no effect on PPI [96], a higher dose (10 mg/kg) disrupted PPI [32], similar to our findings in rats that received acute high-dose THC (20 mg/kg). This suggests the possibility of a dose-dependent effect of THC on PPI. A previous study reported decreased PPI in rats following the direct infusion of CB1R agonists into the hippocampus and PFC [35] and concluded that hippocampus and PFC CB1R activation modulates neural GABA and glutamate release, which affects the activities of PPI-mediating structures like the NAc and ventral tegmental area [35]. This conclusion is consistent with our finding of decreased PFC and dHipp gamma power following acute high-dose THC administration, which may have influenced NAc activity.

Although we report several novel findings, there are a number of limitations that are noteworthy. Firstly, we could only administer the high-dose THC once, since the rats consume the Nutella voluntarily, but do not consume it after exposure to N-THC. Secondly, blood sampling to determine serum THC and 11-OH-THC levels began at the 4-h time-point, just after the AAT and ASR. This was done to ensure that the behavioral effects of the N-THC exposure would not be impacted by the blood sampling, but as a result we may have missed the peak THC and 11-OH-THC levels. Thirdly, although male rats metabolize THC preferentially into 11-COOH-THC, we did not evaluate serum and brain 11-COOH-THC levels. This will be done in future studies. Fourthly, while we chose to use full-spectrum THC oil to more closely mimic human edible cannabis products, we are unable to assess the impact of additional cannabinoids and terpenes in the cannabis oil on the outcomes measured here. Future studies exploring different types of cannabis oils with varying cannabinoid and terpenoid levels may help to clarify these effects. Fifthly, the estrous cycle phase was not assessed in the female rats, which may have impacted the behavioral and electrophysiological effects of edible cannabis. Future studies will be adequately powered to assess the impact of estrous cycle on cannabis poisoning. Lastly, while the model proposed here will be valuable, these findings will need to be extended to younger rats to more closely model cannabis poisoning in children.

In conclusion, edible cannabis-induced poisoning results in decreased gamma power, causes hypolocomotion, hypothermia, and anti-nociception, leads to learning deficits, and impairs PPI. While it is not surprising that cannabis poisoning has neural and behavioral effects, this is the first study to show these effects using edible cannabis, which is increasingly becoming popular among cannabis users and is the most cited cannabis product associated with cannabis poisoning, especially in children and pets. Interestingly, the neural and behavioral effects of edible cannabis-induced poisoning begun after 2 h, which coincides with the period when the peak effects of edible cannabis on cognitive processes were reported. This suggests that administering interventions during this time window may be key to reversing the effects of cannabis poisoning. Moreover, the sex differences in gamma power and serum and brain THC and 11-OH-THC levels following cannabis poisoning suggests sex differences in the effects of THC and its metabolism, which should be considered for the development of effective treatments.

## AUTHOR CONTRIBUTIONS

RQA contributed to the design of the work; acquisition, analysis, interpretation of the data; and drafting and revising the manuscript. HK contributed to the design of the work, acquisition of the data, and revising the manuscript. MAT contributed to the acquisition of the data. AH contributed to the acquisition of the data. YG contributed to the acquisition, analysis, and interpretation of the data. RJ contributed to the acquisition, analysis, and interpretation of the data. KU contributed to the conception of the work, interpretation of the data, and revision of the manuscript. JYK contributed to the conception and design of the work, analysis and interpretation of the data, and the drafting and revision of the manuscript. All authors have approved the final version of the manuscript to be published and agree to be accountable for all aspects of the work in ensuring that questions related to the accuracy and integrity of any part of the work are appropriately investigated and resolved.

---

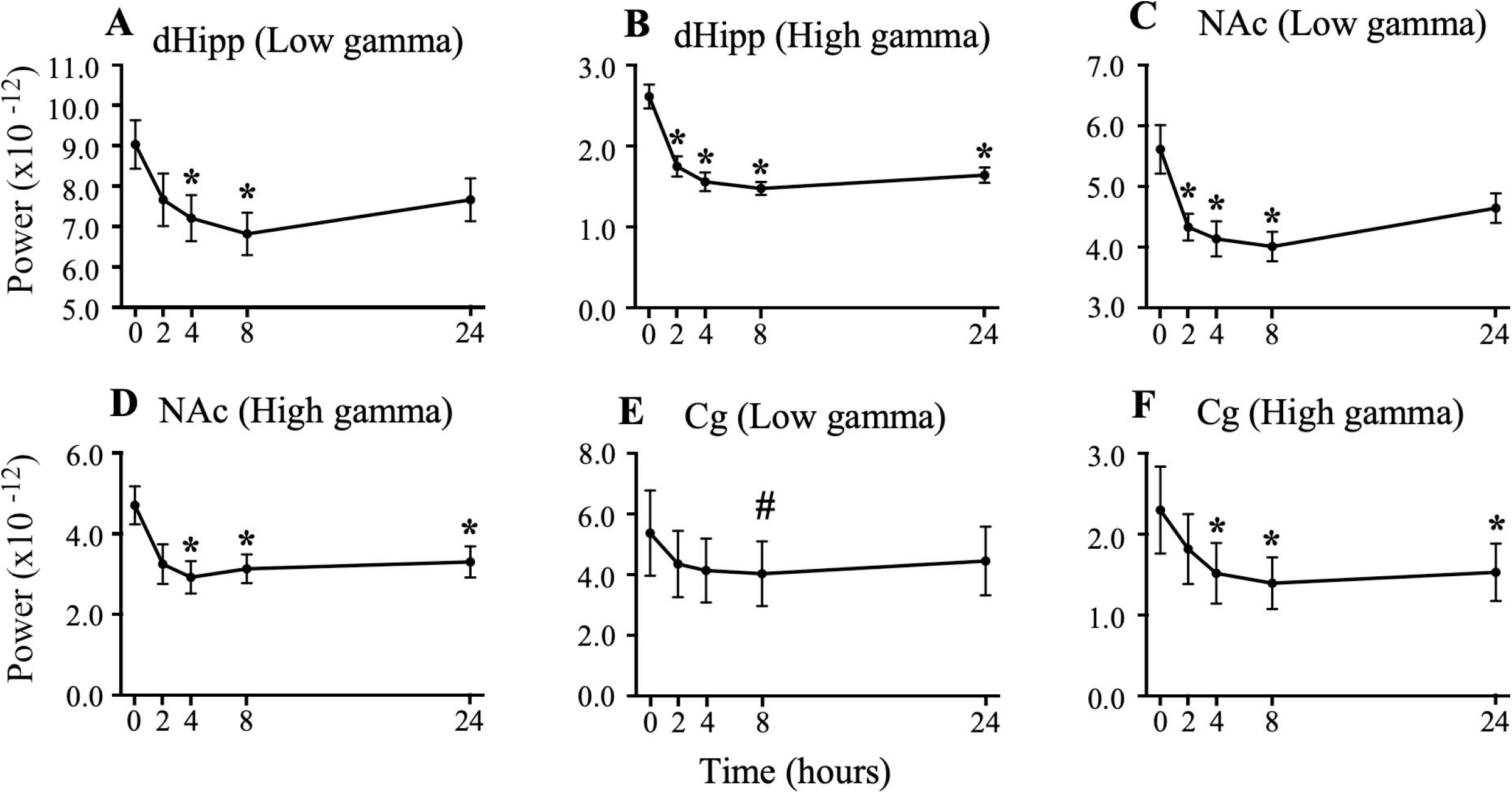

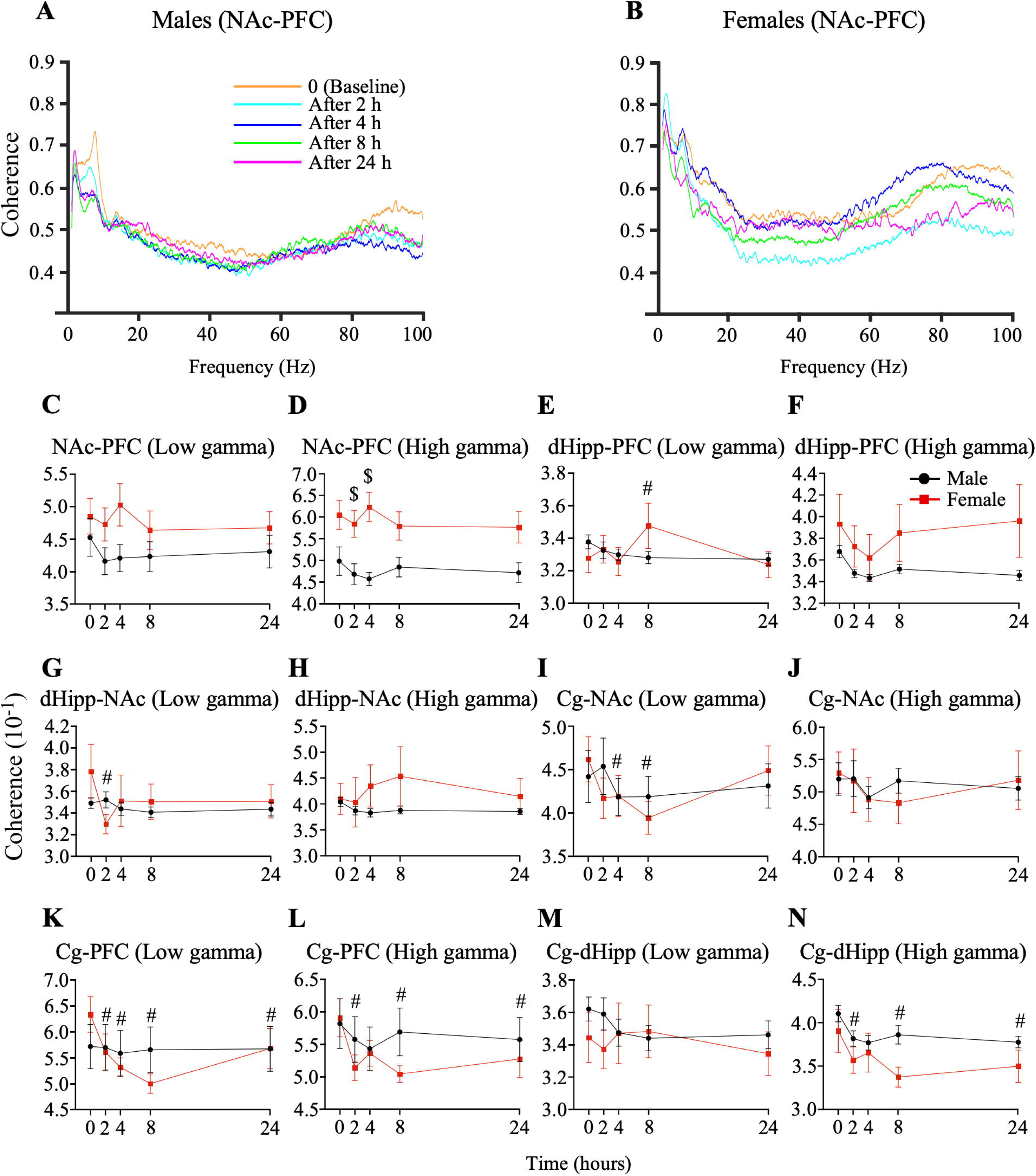

