## Supplemental File for "Neural and behavioral correlates of edible cannabis-induced poisoning: characterizing a novel preclinical model"

Table of content

Supplementary Figure 1: Comparison of low and high gamma power at other time points with baseline in male rats using one-way analysis of variance results

Supplementary Figure 2: Coherence plot for male and female rats after cannabis-edible induced poisoning

Supplementary Table 1. Optimized LC-MS/MS compound parameters for quantitation of 11-OH-THC and THC using PRM mode.

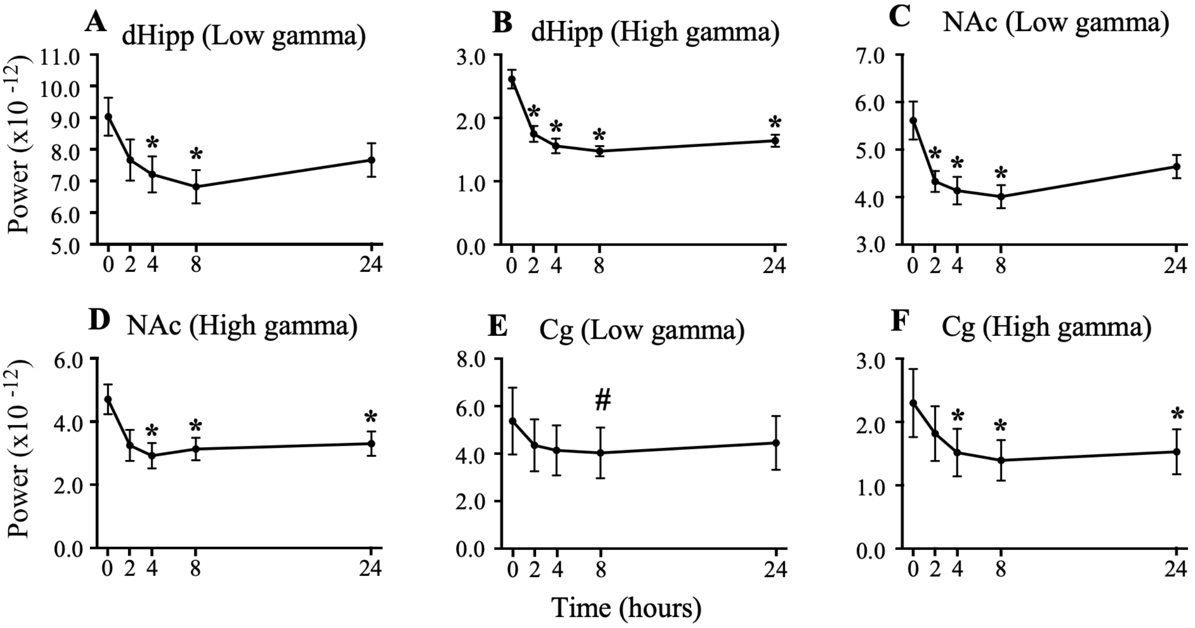


Supplementary Figure 1: Comparison of low and high gamma power at other time points with baseline in male rats using one-way repeated measures analysis of variance. A and B. Graphs comparing low and high gamma power spectral densities, respectively, in the dorsal hippocampus (dHipp) at the 2, 4, 8, and 24-h time-points with that at baseline. C and D. Graphs comparing low and high gamma power spectral densities, respectively, in the cingulate cortex (Cg) at the 2, 4, 8, and 24-h time-points with that at baseline. D and E. Graphs comparing low and high gamma power spectral densities, respectively, in the nucleus accumbens (NAc) at the 2, 4, 8, and 24-h time-points with that at baseline. *: significantly different compared to baseline, #: tending towards significance compared to baseline.

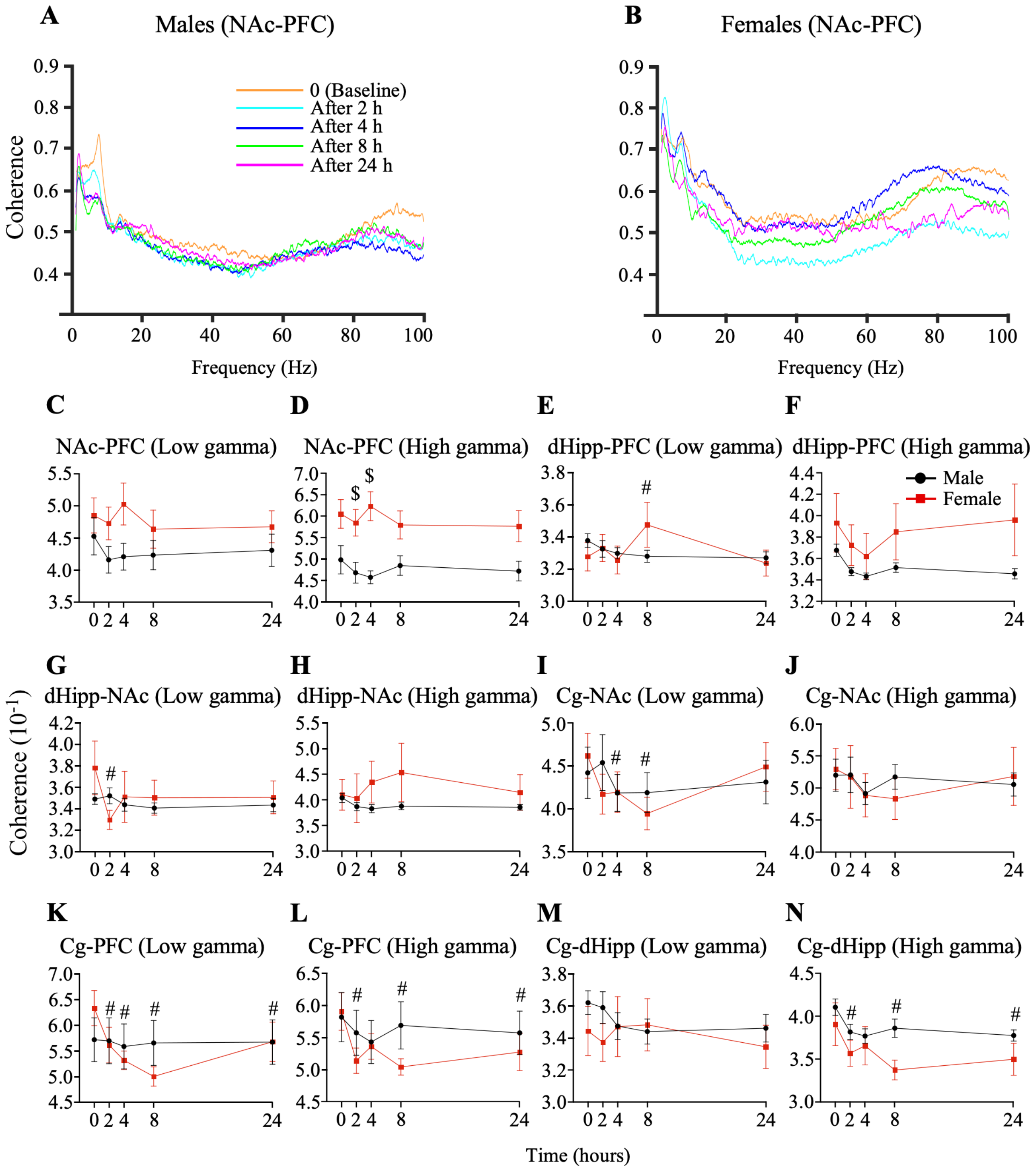


Supplementary Figure 2. Coherence plots for male and female rats after cannabis-edible induced poisoning. Representative coherence plots for male rats (A) and female rats (B) showing the coherence between the nucleus accumbens and prefrontal cortex (NAc-PFC) for each time-point of interest. Orange plot: Baseline, cyan plot: 2-h time-point, blue plot: 4-h time-point, green plot: 8-h time-point, and pink plot: 24-h time-point. C. Graph comparing NAc-PFC low gamma coherence between male and female rats at different time-points. D.

Graph comparing NAc-PFC high gamma coherence between male and female rats at different time-points. E. Graph comparing dorsal hippocampus and prefrontal cortex (dHipp-PFC) low gamma coherence between male and female rats at different time-points. F. Graph comparing dHipp-PFC high gamma coherence between male and female rats at different time-points. G. Graph comparing dorsal hippocampus and nucleus accumbens (dHipp-NAc) low gamma coherence between male and female rats at different time-points. H. Graph comparing dHipp-NAc high gamma coherence between male and female rats at different time-points I. Graph comparing cingulate cortex and nucleus accumbens (Cg-NAc) low gamma coherence between male and female rats at different time-points. J. Graph comparing Cg-NAc high gamma coherence between male and female rats at different time-points. K. Graph comparing cingulate cortex and prefrontal cortex (Cg-PFC) low gamma coherence between male and female rats at different time-points. L. Graph comparing Cg-PFC high gamma coherence between male and female rats at different time-points. M. Graph comparing cingulate cortex and dorsal hippocampus (Cg-dHipp) low gamma coherence between male and female rats at different time-points. N. Graph comparing Cg-dHipp high gamma coherence between male and female rats at different time-points. $: comparison of different time-points between sexes, #: comparison of other time-points with baseline in female rats.

Supplementary Table 1. Optimized LC-MS/MS compound parameters for quantitation of 11-OH-THC and THC using PRM mode.

| Analyte & Internal Standard | Precursor Ion *(m/z)* | CE | Quantitation ion (*m/z)* | Confirming ion (*m/z)* | RT (min) |
|---|---|---|---|---|---|
| 11-OH-THC | 331.23 | 20 | 313.22 | 193.12 | 3.5 |
| 11-OH-THC–D3 | 334.24 | 20 | 316.23 | 196.14 | 3.5 |

| | | | | | |
|---|---|---|---|---|---|
| THC | 315.23 | 25 | 193.12 | 259.17 | 4.5 |
| THC–D3 | 318.25 | 25 | 196.14 | 262.19 | 4.5 |

CE: collision energy, m/z: mass/charge, RT: retention time
